# The cholinergic drug galantamine ameliorates acute and subacute peripheral and brain manifestations of acute respiratory distress syndrome in mice

**DOI:** 10.1101/2025.05.17.654675

**Authors:** Aidan Falvey, Santhoshi P. Palandira, Saher Chaudhry, Aisling Tynan, Fernanda Marciano Consolim-Colombo, Christine N. Metz, Michael Brines, Eric H. Chang, Sangeeta S. Chavan, Kevin J. Tracey, Valentin A. Pavlov

## Abstract

Acute respiratory distress syndrome (ARDS) is a life-threatening form of acute lung injury (ALI), which is a common cause of respiratory failure and high mortality in critically ill patients. Long-term mortality and brain dysfunction have been documented in ARDS patients after hospital discharge. Inflammation plays a key role in ALI/ARDS pathogenesis. Neural cholinergic signaling regulates cytokine responses and inflammation. Here, we studied the effects of galantamine, an approved cholinergic drug (for Alzheimer’s disease) on ALI/ARDS severity and inflammation in mice, using a clinically relevant mouse model induced by intratracheal administration of hydrochloric acid and lipopolysaccharide. Mice were treated 30 mins prior to each insult with vehicle or galantamine (4 mg/kg, i.p.). Galantamine treatment significantly decreased bronchoalveolar lavage (BAL) and serum TNF, IL-1β, and IL-6 levels, as well as BAL total protein and myeloperoxidase and lung histopathology in ALI/ARDS mice. In addition, galantamine improved the functional state of mice with ALI/ARDS during a 10-day monitoring and attenuated lung injury and indices of brain inflammation at 10 days. These findings support further studies utilizing this approved cholinergic drug in therapeutic strategies for ARDS and its subacute sequelae.

## Introduction

Acute respiratory distress syndrome (ARDS) is a life-threatening form of acute lung injury (ALI) and respiratory failure. Globally, ARDS affects approximately 3 million patients annually, accounting for more than 10% of intensive care unit admissions ^1,2^. Mortality of patients with ARDS is estimated at 35% to 46% ^2^. Before 2020, ARDS affected more than 200,000 patients resulting in nearly 75,000 deaths annually in the United States alone ^1,3^. There are also long-term complications in ARDS survivors including respiratory symptoms, as well as physical and mental deterioration for months to years post-initial hospitalization and increased mortality ^4–10^. Brain neurological complications and considerable cognitive impairment including deficiencies in executive function, verbal reasoning, memory, and attention have been reported in survivors of ARDS ^10,11^. The COVID-19 pandemic caused by SARS-COV-2 infection added a novel type of insult leading to ALI/ARDS and long-term functional deterioration, including lung complications, physical and neurological manifestations, and cognitive impairment ^10,12–15^. Despite many years of active research, pharmacological treatments for ARDS are limited and its management is largely based on supportive care, including lung-protective mechanical ventilation ^1^.

ALI/ARDS occurs in the context of various pulmonary complications, including pneumonia, toxic inhalation, lung contusion, aspiration, injurious ventilation, and non-pulmonary insults such as sepsis, pancreatitis, trauma, and burns. These insults lead to acute hypoxemia and non-hydrostatic pulmonary edema mediated through epithelium, alveolar macrophages, vascular endothelium alterations, and inflammatory lung injury with the release of pro-inflammatory cytokines ^1,10,16^. The exacerbated release of pro-inflammatory cytokines such as tumor necrosis factor (TNF) and interleukin-6 (IL-6) from mononuclear cells during ALI/ARDS and their increased circulating levels mediates systemic inflammation which plays a major role in morbidity and mortality in patients with ARDS ^16–22^. It has been acknowledged that in addition to initial phases, providing insights into ALI/ARDS at subacute stages using appropriate animal models may inform further developments in the pursuit of new treatments ^23,24^. However, investigations in rodent models continue to focus on ALI/ARDS pathobiology within the first 1-4 days after onset and few studies have examined the acute and subacute ALI/ARDS phases ^24–26^.

It has been acknowledged that in addition to initial phases, providing insights into ALI/ARDS at subacute stages using appropriate animal models may inform further developments in the pursuit of new treatments ^23,24^. However, investigations in rodent models continue to focus on ALI/ARDS pathobiology within the first 1-4 days after onset and only a few studies have examined acute and subacute ALI/ARDS phases ^24–26^.

Brain and vagus nerve cholinergic mechanisms play a critical role in controlling pro-inflammatory cytokine responses and inflammation ^27–34^. Cholinergic activation can be achieved using galantamine, an acetylcholinesterase (AChE) inhibitor approved for treating Alzheimer’s disease ^35^. We and others have shown that galantamine activates brain cholinergic anti-inflammatory signaling and efferent vagus nerve activity leading to suppression of pro-inflammatory cytokines in endotoxemia, colitis, obesity, and other inflammatory conditions ^35–39^. These studies provided a rationale for successful clinical exploration demonstrating the anti-inflammatory and beneficial cardiometabolic effects of galantamine in people with the metabolic syndrome ^40,41^.

In this study, we used a clinically relevant mouse model of ALI/ARDS ^26^ to examine the effects of treatment with galantamine on the severity of ALI/ARDS and inflammation in acute (30 h) and subacute (10 days) settings. Our results demonstrate that galantamine alleviates lung tissue injury and local and systemic inflammation at 30 h, as well as lung injury and neuroinflammation at 10 days. These findings show that this approved drug should be further examined as a potential treatment for ARDS.

## Results

### Galantamine suppresses BAL and circulating pro-inflammatory cytokine levels in mice with ALI/ARDS

Increased lung and circulating pro-inflammatory cytokines mediating local and systemic inflammation are a hallmark in patients with ALI/ARDS and cause broader pathogenesis and organ dysfunction ^16–22^. We used a model of ALI/ARDS generated by intratracheal (i.t.) administration of hydrochloric acid (HCl, 0.1N, 2 ml/kg) and lipopolysaccharide (LPS, endotoxin, 10 mg/kg) 24h later, as depicted in **Figure 1A**. As previously highlighted, this model provides a clinically relevant scenario of lung and systemic inflammation, pronounced lung injury, and brain neurological complications at subacute stages ^26^. We first examined the effects of galantamine (4 mg/kg, i.p.) injected 30 mins prior to HCl and endotoxin (**Figure 1A**) on bronchoalveolar (BAL) and serum cytokine levels in mice with ALI/ARDS at 30h and the effects of galantamine on these levels. As shown in **Figure 1B**, BAL TNF, IL-6, and IL-1β levels were significantly elevated in ALI/ARDS mice compared with control mice. Galantamine treatment significantly decreased BAL IL-6 levels in ALI/ARDS mice to levels no different than those in controls and significantly suppressed BAL IL-1β levels. Serum TNF, IL-6, and IL-1β levels were also significantly higher in mice with ALI/ARDS compared with controls (**Figure 1C).** Of note, galantamine treatment significantly decreased all these cytokines in mice with ALI/ARDS to levels, that were not statistically different from controls. Collectively these results indicate significant anti-inflammatory effects of cholinergic activation by galantamine in murine ALI/ARDS.

**Figure 1.**
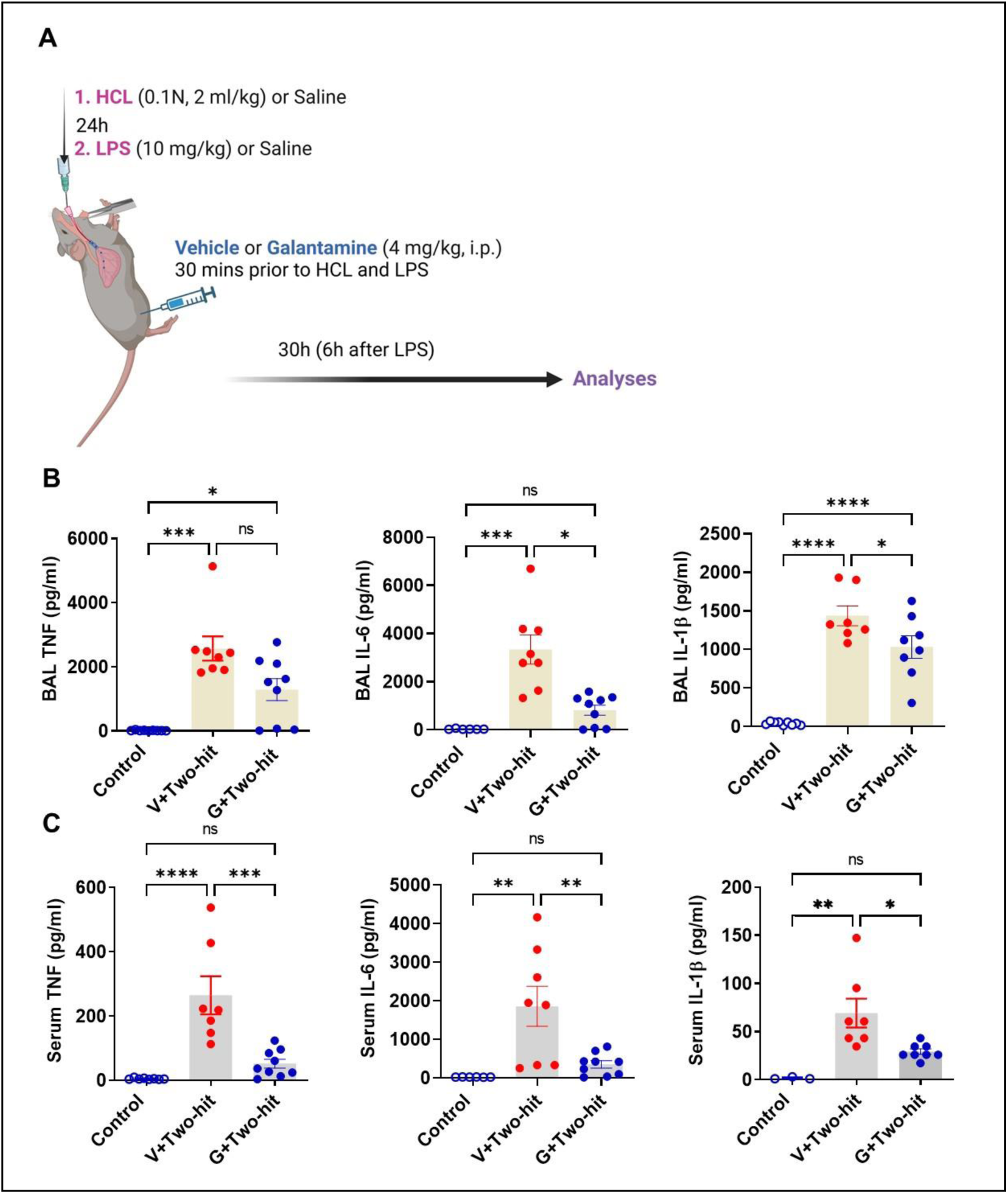
Galantamine treatment suppresses pro-inflammatory cytokine levels in mice with ALI/ARDS. **(A**) Schematic depiction of the experimental design. Anesthetized mice were administered with HCL and LPS (i.t.) 24h apart and treated 30 mins prior to each insult with galantamine (4 mg/kg) or vehicle administered i.p. Another (control) group of anesthetized mice was subjected to the same experimental procedure but administered vehicle (saline). Mice were euthanized at 30h and blood, bronchoalveolar lavage (BAL), lung and brain (at 10 days) were collected and processed for analysis. (**B**) BAL TNF levels are significantly higher in mice with ALI/ARDS treated with vehicle (V) (***P=0.0002; Kruskal Wallis test, Dunn’s multiple comparisons test) and in mice with ALI/ARDS treated with galantamine (G) (*P=0.0486) compared with control mice. BAL IL-6 levels are significantly higher in mice with ALI/ARDS treated with vehicle compared with control mice (***P=0.0002; Kruskal Wallis test, Dunn’s multiple comparisons test) and galantamine significantly decreases these levels (*P=0.0226). BAL IL-1β levels are significantly higher in mice with ALI/ARDS treated with vehicle (****P<0.0001; one-way ANOVA, Tukey’s multiple comparisons test) and in mice with ALI/ARDS treated with galantamine (****P<0.0001) compared with control mice. Galantamine significantly decreases IL-1β levels in mice with ALI/ARDS compared with mice treated with vehicle (*P=0.0422). (**C**) Serum TNF levels are significantly higher in mice with ALI/ARDS treated with vehicle than control mice (****P<0.0001; one-way ANOVA, Tukey’s multiple comparisons test) and galantamine significantly decreases these values (***P=0.0003). Serum IL-6 levels are significantly higher in mice with ALI/ARDS treated with vehicle than control mice (**P=0.0031; one-way ANOVA, Tukey’s multiple comparisons test) and galantamine significantly decreased these values (**P=0.0065). Serum IL-1β levels are significantly higher in mice with ALI/ARDS treated with vehicle than control mice (**P=0.017; Kruskal Wallis test, Dunn’s multiple comparisons test) and galantamine significantly decreases these values (*P=0.0430). Data are represented as individual mouse data points with mean ± SEM. See Materials and Methods for details.

### Galantamine treatment alleviates acute lung injury in mice

We also studied the effects of galantamine on BAL markers of lung injury and lung tissue damage evaluated histologically at 30h after onset. As shown in **Figure 2 A, B**, at 30h the levels of total protein and myeloid peroxidase (MPO) in the BAL were significantly elevated in mice subjected to the two-hit ALI/ARDS and injected with vehicle compared with control mice, processed similarly, but administered i.t. with saline. Galantamine significantly decreased BAL total protein and BAL MPO at 30h to levels that were not statistically different compared with control mice **(Figure 2 A, B).** Histological examination of the lung in the three groups of mice revealed a greater presence of neutrophils in the alveolar space and cell walls, as well as instances of hyaline membranes, and alveolar wall thickening in ALI/ARDS mice treated with vehicle compared with controls and ARDS mice treated with galantamine (**Figure 2 C**). Whole lung sections are shown in **Supplementary Figure 1.** Quantitative evaluation implementing a scoring system as per The American Thoracic Society ^42^ demonstrated higher lung injury scores in ALI/ARDS mice compared with controls and significantly decreased scores in ALI/ARDS mice treated with galantamine (**Figure 2 D**). These results show that galantamine ameliorates acute lung injury in mice.

**Figure 2.**
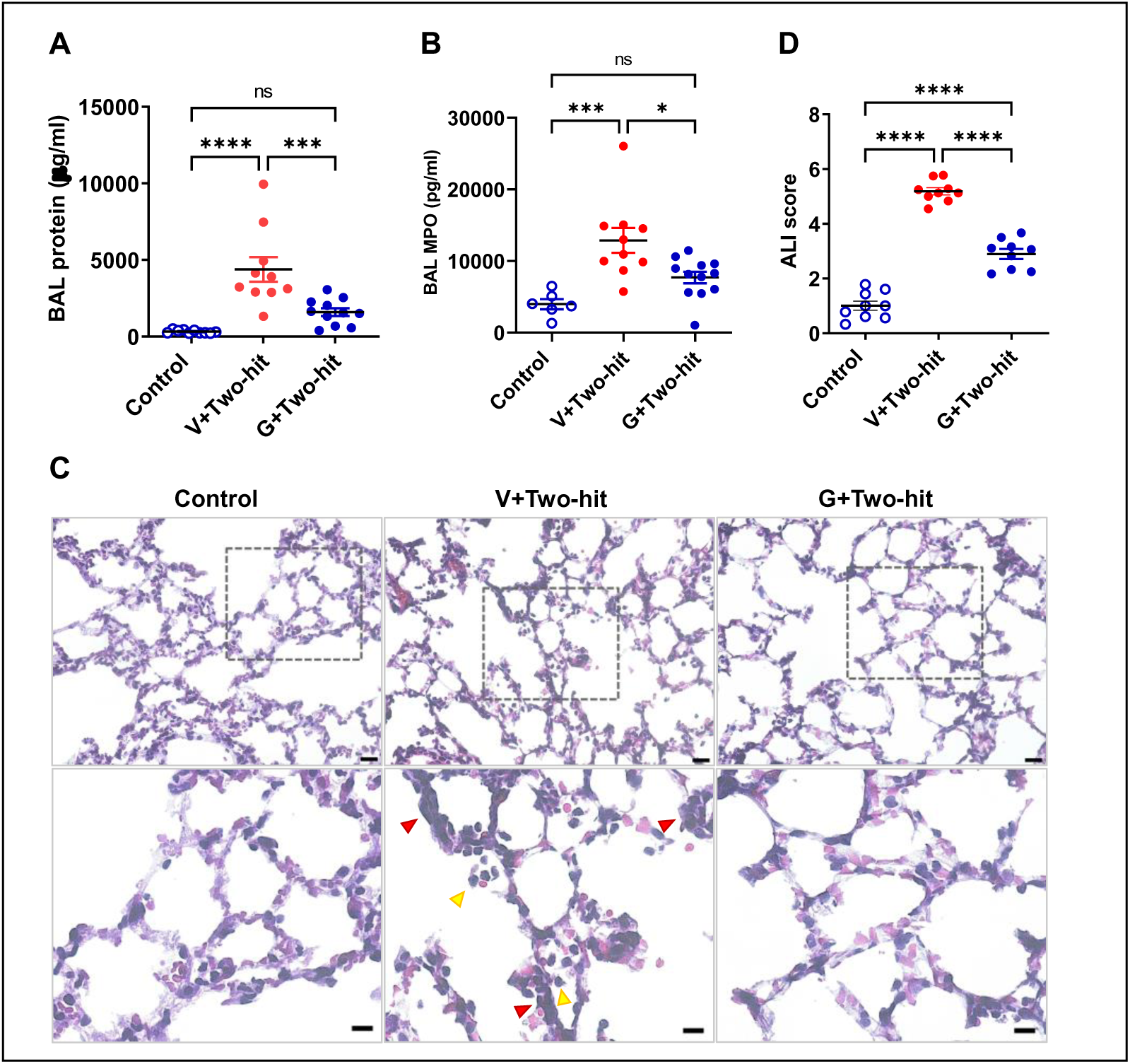
Galantamine treatment ameliorates markers of acute lung damage in mice with ALI/ARDS. (**A**) At 30h BAL total protein levels are significantly higher in mice with ALI/ARDS treated with vehicle (Veh) (****P<0.0001; one-way ANOVA, Tukey’s multiple comparisons test) compared with control mice and galantamine treatment significantly decreases BAL total protein in ALI/ARDS mice (**P=0.0007) to levels no different than controls. (**B**) BAL MPO levels are significantly higher in mice with ALI/ARDS treated with vehicle than control mice (***P<0.0005; one-way ANOVA, Tukey’s multiple comparisons test) and galantamine significantly decreases these values (***P=0.00125) to levels no different than controls. (**(C)** Representative H&E images of mouse lung tissue sections show the degree of ALI injury in the two-hit model, including neutrophil infiltration in alveolar space (yellow arrowheads) and interstitial space (red arrowheads). Top row images, scale bar = 20 µm; bottom row images, scale bar = 10 µm. **(D**) The acute lung injury score in mice with ALI/ARDS treated with vehicle (****P<0.0001; one-way ANOVA, Tukey’s multiple comparisons test) and those treated with galantamine (****P<0.0001) is significantly higher compared with controls. Galantamine treatment of mice with ALI/ARDS significantly decreases the score compared with vehicle treatment (****P<0.0001). Each individual point represents an average of 5 sections that were scored. Data are represented as individual mouse data points with mean ± SEM. See Materials and Methods for details.

### Galantamine improves general functional activity in mice with ALI/ARDS

We next studied functional alterations in the three groups of mice implementing a 10-day monitoring protocol and utilizing a previously established non-invasive scoring system ^43^. This system, originally developed for assessing mice with sepsis is based on seven clinical variables, including appearance, level of consciousness, activity, response to stimulus, eye opening, respiratory rate, and quality of respirations ^43^. Because alterations in all these variables are present in mice with ALI/ARDS, we reasoned that the use of this scoring system would be appropriate. As shown in **Figure 3A**, the functional/illness score of mice subjected to the two-hit (i.t.) ALI/ARDS and treated with vehicle was significantly worsened compared with control mice. The illness severity was significantly ameliorated in ALI/ARDS mice treated with galantamine. We also monitored body weight of the three groups of mice for 10 days. As shown in **Figure 3B** a significant overall body weight loss was observed in mice with ALI/ARDS treated with vehicle compared with control mice. However, no statistically significant difference was found between control mice and mice subjected to ALI/ARDS and treated with galantamine. These results indicate that the beneficial effects of galantamine on acute lung injury and inflammation are associated with alleviated illness severity and improved overall functional activity, with lack of significant body weight loss.

**Figure 3.**
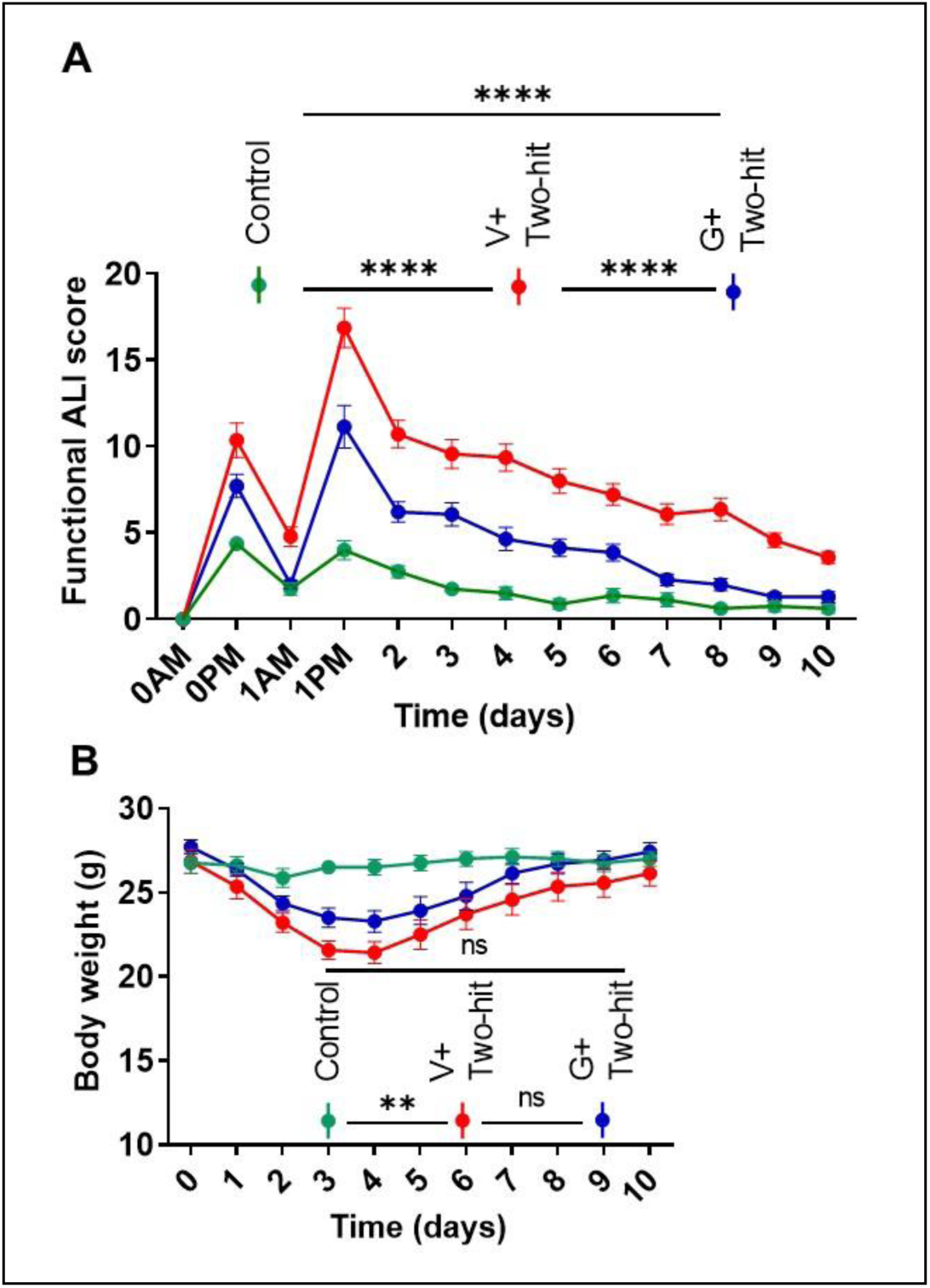
Galantamine treatment improves the functional activity of mice with ALI/ARDS with no effect on body weight. **(A)** Mice with ALI/ARDS treated with vehicle (n=14) or galantamine (n=14) have impaired functional activity indicated by high illness scores compared with controls (n=8) (***P<0.0001; repeated measures two-way ANOVA, Tukey’s multiple comparisons test) during 10-day monitoring. Galantamine treatment improves the functional activity of mice with AIL/ARDS reflecting a significantly lower illness score compared with vehicle treatment (***P<0.0001). (**B**) The overall body weight of mice with ALI/ARDS treated with vehicle (n=14) is significantly lower compared with controls (n=8) (**P=0.0058; repeated measures two-way ANOVA, Tukey’s multiple comparisons test) and not significantly different than ALI/ARDS mice treated with galantamine during 10-day monitoring. See Materials and Methods for details.

### Galantamine alleviates subacute lung injury in mice with ALI/ARDS

We next examined lung injury indices in the three groups of mice after 10 days. BAL total protein levels were significantly increased in mice with ALI/ARDS compared with control mice (**Figure 4A**). Similarly, BAL MPO in ALI/ARDS mice was significantly elevated compared with controls (**Figure 4B**). However, there were no statistically significant differences in BAL total protein and MPO between control mice and mice with ALI/ARDS treated with galantamine (**Figure 4A, B**). There also were overall trends to lower BAL total protein and BAL MPO in ALI/ARDS mice treated with galantamine compared with ALI/ARDS mice treated with vehicle. In addition to these markers of local tissue injury, we measured BAL and circulating TNF levels in mice in the three groups at 10 days after onset. BAL TNF levels in mice with ALI/ARDS were very low (as compared to those determined at 30h, **Figure 2A**) and no different than TNF levels in control mice or in ALI/ARDS mice treated with galantamine (**Supplementary Figure 2A)**. Similarly, we determined very low serum TNF levels in ALI/ARDS mice and no significant differences comparing the three groups of mice (**Supplementary Figure 2B**). We also measured BAL levels of the anti-inflammatory cytokine IL-10 ^44^ with a recognised role in decreasing lung inflammation ^45^. Low BAL IL-10 levels were detected with no significant difference between the three groups of mice (**Supplementary Figure 2C**). As shown on representative images (**Figure 4C),** histological evaluation of the lung across the three groups of mice at 10 days revealed persistent lung damage, including prominent infiltration of neutrophils in the alveolar and interstitial spaces, alveolar wall thickening, and sparse proteinaceous in mice subjected to ALI/ARDS compared with controls and ALI/ARDS mice treated with galantamine. Whole lung sections are shown in **Supplementary Figure 3.** Accordingly, the lung injury score was significantly higher in mice with ALI/ARDS treated with vehicle compared with controls and galantamine treatment of ALI/ARDS mice significantly lowered the score to levels that were not statistically different compared with control mice (P=0.06, One-way ANOVA, Kruskal Wallis) (**Figure 4D**). Together these results show that galantamine attenuates features of persistent subacute (at 10 days) lung injury in mice with ALI/ARDS with no significant effect on low pro-inflammatory and anti-inflammatory cytokine levels at this stage (where the inflammatory state has subsided).

**Figure 4.**
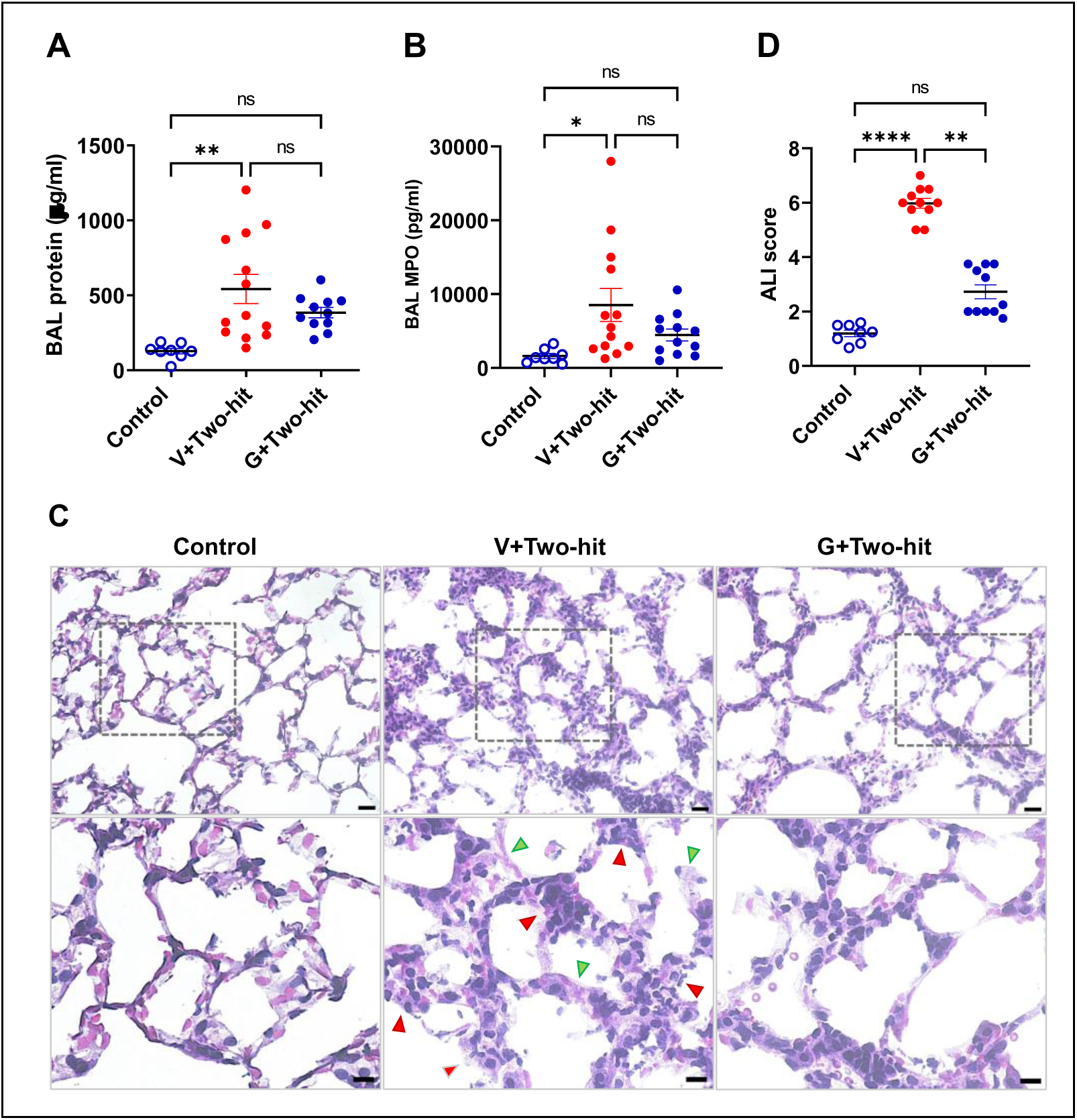
Galantamine treatment attenuates indices of subacute ALI/ARDS in mice. (**A**) BAL total protein levels are significantly higher in mice with ALI/ARDS treated with vehicle (Veh) (****P=0.0014; one-way ANOVA, Tukey’s multiple comparisons test) compared with control mice and not significantly different than ALI/ARDS mice treated with galantamine. (**B**) BAL MPO levels are significantly higher in mice with ALI/ARDS treated with vehicle than control mice (***P=0.0199; one-way ANOVA, Tukey’s multiple comparisons test) and not significantly different than ALI/ARDS mice treated with galantamine. (**C**) Representative H&E images of mouse lung tissue sections show the degree of ALI injury in the two-hit model after 10 days, including neutrophil infiltration in interstitial space (red arrowheads) and thickening of hyaline membranes (green arrowheads). Top row images, scale bar = 20 µm; bottom row images, scale bar = 10 µm. (**D**) The lung injury score in mice with ALI/ARDS treated with vehicle is significantly higher compared with controls (****P<0.0001; Kruskal Wallis test, Dunn’s multiple comparisons test). Galantamine treatment of mice with ALI/ARDS significantly decreases the score compared with vehicle treatment (**P<0.0099). Each individual point represents an average of 5 sections that were scored. Data are represented as individual mouse data points with mean ± SEM. See Materials and Methods for details.

### Galantamine reduces microglial accumulation in the hippocampus of mice with ALI/ARDS

In addition to lung and systemic pathobiological sequalae, persistent cognitive impairment has been documented in ARDS survivors after hospital discharge ^6^. Behavioral alterations and impaired spatial learning and memory have been previously associated with hippocampal alterations at subacute stages in a two-hit ALI/ARDS murine model ^26^. Both clinical and animal studies have demonstrated an association between brain inflammation (neuroinflammation) and microglial alterations, cognitive impairment, and hippocampal neuronal dysfunction ^46^. Accordingly, we next studied whether there were microglial alterations in the hippocampus in the three groups of mice. Using IBA1 immunolabeling, we observed significantly higher numbers of microglia in mice with ALI/ARDS compared with controls. The number of microglia in the hippocampus of ALI/ARDS mice treated with galantamine was significantly lower compared with the ALI/ARDS mice treated with vehicle and not statistically different than control mice (**Figure 5A and 5B).** Additional analysis of hippocampal microglial ramification implementing a previously utilized methodology ^47,48^ as shown in **Supplementary Figure 4** demonstrated significantly increased values in mice with ALI/ARDS compared with controls (**Figure 5C**). Galantamine treatment of mice with ALI/ARDS significantly reduced these values to levels, which were not statistically different than controls (**Figure 5C**). These results indicate increased inflammation in the hippocampus in mice with ALI/ARDS at a subacute (10 day) stage and the counteracting anti-inflammatory effect of galantamine treatment.

**Figure 5.**
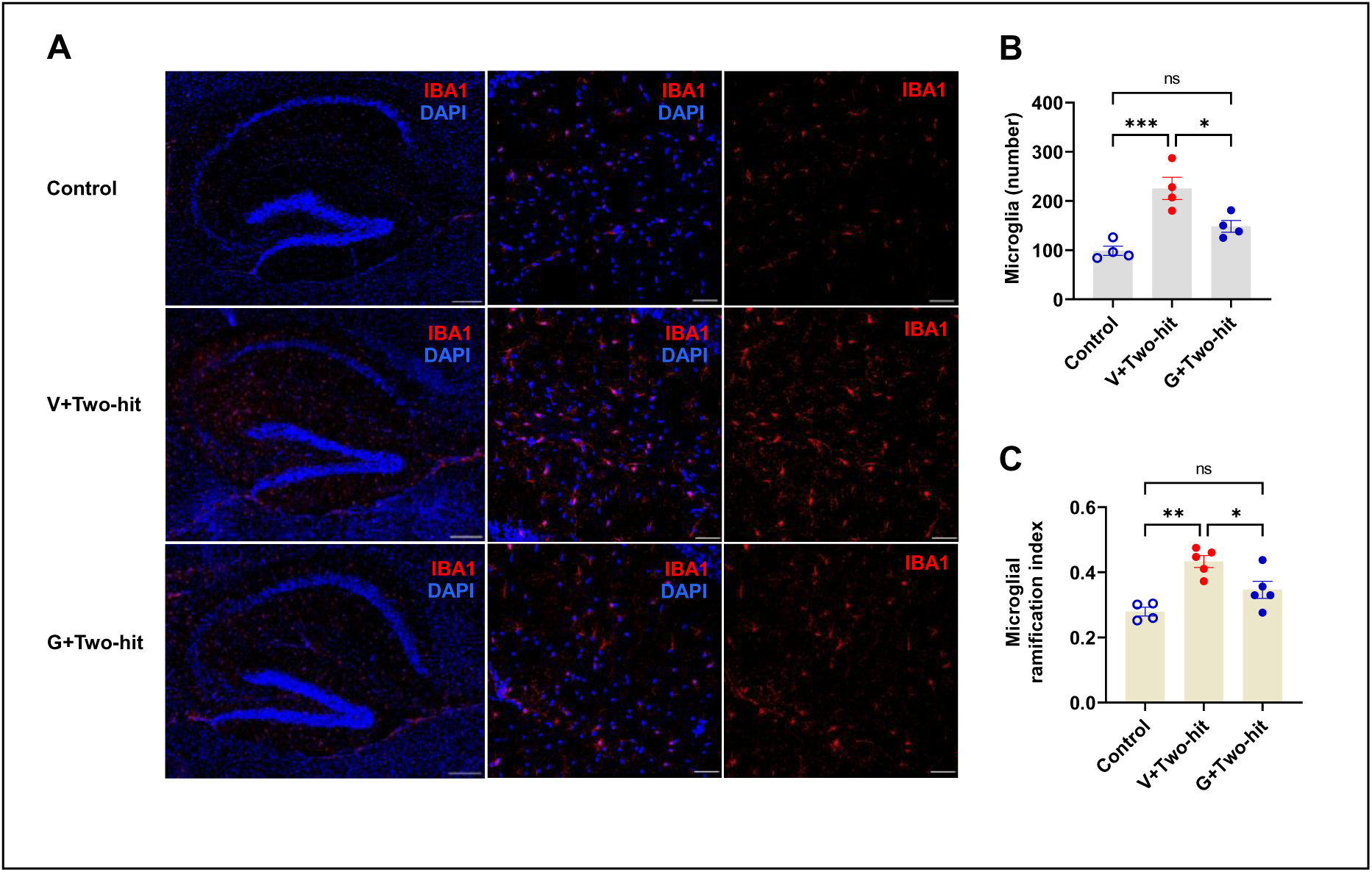
Galantamine treatment decreases microglial accumulation in the hippocampus of mice with subacute ALI/ARDS. **(A)** Representative images of IBA1 staining of brain (hippocampus) shown in wide tiles (Scale bar = 200 µm) and with 20x magnification (Scale bar = 50 µm in the three groups of mice. **(B)** The number of microglia in the hippocampus of mice with ALI/ARDS treated with vehicle is significantly higher compared with controls (***P=0.0008; one-way ANOVA, Tukey’s multiple comparisons test). Galantamine treatment of mice with ALI/ARDS significantly decreases microglial accumulation in the hippocampus compared with vehicle treatment (*P=0.0181). (**C**) The ramification index of microglia in the hippocampus of mice with ALI/ARDS treated with vehicle is significantly higher compared with controls (**P=0.001; one-way ANOVA, Tukey’s multiple comparisons test). Galantamine treatment of mice with ALI/ARDS significantly decreases the microglial ramification index in the hippocampus compared with vehicle treatment (*P=0.029). Data are represented as individual mouse data points with mean ± SEM. See Materials and Methods for details.

## Discussion

ALI/ARDS is a common life-threatening disease with high in-hospital mortality. ARDS mortality extends beyond the acute phase in parallel with considerable physical, mental, and cognitive deterioration for years after hospital discharge. This dictates the need for finding better treatments with good safety profiles. Local lung and systemic inflammation with increased pro-inflammatory cytokines play a major role in ALI/ARDS pathogenesis. Here we show that galantamine, an AChE inhibitor and a cholinergic compound with previously established anti-inflammatory activity, alleviates murine ALI/ARDS and attenuates local (BAL) and systemic pro-inflammatory cytokine levels. In addition, galantamine treatment improves the functional activity of mice with ALI/ARDS monitored for 10 days, ameliorates lung injury, and decreases hippocampal neuroinflammation at subacute stages of the condition.

Pro-inflammatory cytokines, including TNF, IL-6, and IL-1β are validated markers of ongoing cellular injury and their increased BAL levels have been associated with increased morbidity and mortality in patients with ALI/ARDS ^19,49^. These increases reflect simultaneous elevation in circulating pro-inflammatory cytokine levels driving further pathogenesis ^49^. In our study, at 30h galantamine inhibited BAL and systemic pro-inflammatory cytokines such as TNF, IL-6, and IL-1β - in many instances to levels not significantly different than those in control mice. BAL total protein is a recognized marker of vascular/capillary leakage and BAL myeloperoxidase (MPO), released by activated neutrophils, is a validated indicator of accumulation of activated neutrophils within the lung airspaces ^42^. At 30h, the significantly lower levels of BAL total protein and MPO in ALI/ARDS mice treated with galantamine are consistent with a beneficial effect of this cholinergic drug on these processes, which play a key role in ALI pathogenesis. In line with these observations, histological assessment of the lung revealed that galantamine also significantly alleviated ALI.

Long-term mortality and morbidity in patients surviving ARDS remains a significant medical problem. While abundant insights into the initial inflammatory injury have been provided, the subacute consequences of ALI/ARDS in murine models have not been adequately evaluated. We found that the effects of galantamine on ALI were associated with significantly reduced illness score in mice with ALI/ARDS compared to vehicle-treated mice during the 10-day monitoring. Histological analysis revealed the presence of significant lung injury and notably an increased alveolar wall thickness in ALI/ARDS mice at 10 days following the initial insult. This finding is consistent with the persistent lung injury documented in patients surviving ARDS ^13,50^. Importantly, galantamine treatment considerably and significantly prevented/attenuated these alterations.

Histological findings in mice with ALI/ARDS were in line with significantly increased BAL total protein and MPO levels, albeit not to levels observed in acute settings (at 30h). Of note, these increases in galantamine treated animals compared with control mice were not significant. We also found that the persistent lung injury at subacute stages was not paralleled with significant alterations in BAL and serum pro-inflammatory cytokine levels.

In the brain, we found a significantly higher density of hippocampal microglia with increased ramifications in mice with ALI/ARDS, with galantamine decreasing these to control levels. Previous studies have indicated the impact of increased peripheral pro-inflammatory cytokines as well as inflammatory and metabolic derangements on the brain associated with microglial alterations indicative of neuroinflammation ^51–53^. In addition to being a key component of brain immunity, microglia interact with neurons to maintain neuronal integrity and function ^54,55^. Microglial activation and higher numbers drive brain inflammation with a deleterious effect on the neurons ^29,56,57^. The hippocampus is a brain area in which microglial alterations have been documented during aberrant systemic inflammation and increased circulatory levels of IL-6 and other cytokines ^56,57^. Hence, while no difference in pro-inflammatory cytokines were detected at 10 days, it is reasonable to conclude that increased circulating pro-inflammatory cytokine levels at early stages played a causative role in the brain microglial alterations. Furthermore, this neuroinflammation in the hippocampus may contribute to the previously shown cognitive impairment at these late stages of ALI/ARDS in this murine model ^26^. Our results indicating that galantamine alleviates inflammation in the hippocampus are consistent with previous findings that galantamine decreases neuroinflammation in other mouse models of inflammatory conditions ^35^. Galantamine is an AChE inhibitor that increases brain cholinergic signaling which plays an important role in cognition^58,59^. This cholinergic drug is approved for the symptomatic treatment of cognitive impairment in Alzheimer’s disease^59,60^ with a sustained (36 months) effect ^61^.

Previous studies have revealed the robust anti-inflammatory efficacy of galantamine ^35,40,41^ and its beneficial metabolic and cardioprotective effects ^35,40,62–65^. In our study we used a galantamine dose of 4 mg/kg, with previously demonstrated anti-inflammatory and beneficial metabolic effects in mice with endotoxemia ^36^, colitis ^37^, obesity and metabolic syndrome ^39^, pancreatitis ^66^ and other conditions ^35,67^. This dose provides an approximate human equivalent dose calculated allometrically ^68^ to be 0.33 mg/kg body weight respectively, which is within the dosing range of humans receiving treatment for Alzheimer’s disease (0.13 to 0.40 mg/kg body weight). Therefore, our findings are of significant interest for direct translational studies in patients with ARDS. Such investigations can be significantly facilitated by the abundant available information indicating a favorable safety drug profile; in addition to Alzheimer’s disease galantamine has been clinically utilized in many other populations, including children with autism with no notable adverse events^35,69,70^. In addition, galantamine has been successfully used in the treatment of muscle weakness ^35^. Persistent muscle weakness has been documented as a key component of physical deterioration in ARDS survivors for years after acute hospital care, severely worsening their mobility and quality of life ^15,71^. Therefore, it would be of interest to consider galantamine treatment in this context as well.

## Conclusions

Our results indicate that galantamine administered in mice with ALI/ARDS significantly alleviates local (BAL) and circulating cytokine responses and attenuates acute lung damage at early stages (30h). We also show that galantamine improves the overall functional state, attenuates lung injury, and ameliorates neuroinflammatory alterations at subacute stages (10 days) of murine ALI/ARDS. While it is recognized that the long-term ARDS sequelae can be prevented or treated at earliest stages of hospital care ^15^, no efficient targeted pharmacological treatments are available. These findings provide a rationale for studying galantamine, an approved cholinergic drug, in the management of acute and subacute (long-term) ALI/ARDS.

## Materials and Methods

### Animals

The Institutional Animal Care and Use Committee (IACUC) and the Institutional Biosafety Committee of the Feinstein Institutes for Medical Research, Northwell Health, Manhasset, NY approved all procedures described. All procedures were conducted in accordance with NIH guidelines. The study was performed and reported in accordance with ARRIVE guidelines (https://arriveguidelines.org).

Male, C57BL/6 mice were purchased from Jackson Laboratories. All mice were given access to food and water *ad libitum* and maintained on a 12h light and dark schedule at 25°C and fed a standard Purina rodent chow diet (<u>Rodents - Feed and nutrition products | Purina</u> <u>(multipurina.ca)</u>). Mice were allowed to acclimate to the environment for two weeks before being used in experiments. Experiments were conducted on male mice aged 10-12 weeks old.

### Two-hit acute lung injury and treatment with galantamine

To induce lung injury, we followed a previously described procedure ^26^. Mice were anesthetized using ketamine/xylazine (75-100 mg/kg/5-10mg/kg, i.p.) and mounted on an intubation platform (KentScientific). Mice were hung from their front incisors, exposing, and lengthening the trachea. An optical fibre, with an attached safety cap of a catheter needle, was used to locate the trachea and the safety cap was placed into the trachea. Location was verified by attaching a saline filled syringe to the safety cap and observing an apparent rhythmic gas transfer. A catheter was then inserted into the trachea via the safety cap and hydrochloric acid HCl (0.2 N, 2 ml/kg) was slowly instilled. Animals were allowed to recover in a heated cage. 24 h later LPS (10 mg/kg) intratracheal (i.t.) instillation was performed as described above. Another group of control mice was similarly anesthetised and subjected to i.t. saline (instead of HCL and LPS administration). Animals were intraperitoneally (i.p.) injected with vehicle (saline) or galantamine (4 mg/kg. i.p.) 30 mins prior to HCL and LPS i.t. administration. The animals were allowed to recover and 6 h after LPS administration (30h after onset - HCL administration) they were euthanised. Other cohorts of mice were euthanised at 10 days after onset. Animals were euthanized by CO2 asphyxiation. Blood was collected through cardiac puncture. The left lung lobe was excised for histology and bronchoalveolar lavage (BAL) fluid was collected from the right lung. Brains (in the 10-day experiments) were collected) and passively perfused for 72 h in 4% PFA prior to transfer to 30% sucrose and allowed to equilibrate prior to processing for tissue sectioning.

### BAL fluid collection and analyses

Following blood collection via cardiac puncture, the ribcage was fully removed, and the lungs were exposed and the skin/muscle layer covering the trachea was additionally removed. The trachea was pierced with a catheter needle and a guiding syringe was left in the trachea and sutured in place. Three distinct rounds of 0.1 ml + 0.5 ml cold PBS were flushed into and back out of the right lung to collect BAL fluid, samples were frozen and stored until use.

### Lung histochemistry evaluation

Lungs were processed for H&E staining following a previously published method ^72^. In brief, the lungs and trachea were removed, a syringe containing 4% paraformaldehyde was placed into the trachea and the lungs were re-inflated until they returned to their physiological size. Subsequently, the lungs were placed in 4% paraformaldehyde for passive tissue fixing and then 48 h in 30% sucrose. The left lung was covered with OCT, frozen and then stored at -80 degrees Celsius prior to sectioning on a cryostat at 10 microns. Sections of the left lung (10 microns) were stained with H&E following a standard procedure ^72^. Lung sections were histologically evaluated using a scoring system established by the American Thoracic Society (ATS), which yields an acute lung injury (ALI) score ^42^. The scoring system consists of five variables that are scored from 0-2 depending on the lung injury severity: neutrophils in the alveolar space; neutrophils in the interstitial space; hyaline membranes; proteinaceous debris filling the airspaces and alveolar septal thickening. The summative score for each scoring criteria across the lung section was recorded, with a higher ALI score reflecting greater severity of acute lung injury.

### Analyte measurement

BAL total protein was determined using a Bradford analysis kit. MPO activity was determined using (R&D Systems DY3667). Blood was withdrawn and centrifuged for serum analysis. BAL fluid and serum were analyzed for cytokine levels using standard ELISA kits: TNF (88-7324: Invitrogen); IL-6 (R&D Systems DY406); IL-1β (R&D Systems DY401) and IL-10 (R&D Systems DY417). BAL and serum cytokines were expressed as picogram per milliliter of BAL or serum collected. All BAL analyte values were additionally standardized to the volume of BAL fluid collected.

### Functional assessment

Groups of mice were weighted and evaluated using a scoring system involving variables, such as fur quality, consciousness, ambulation, response to stimuli, openness of the eyes, respiration rate and quality, and providing scores from low (0) – high severity (4) ^43^. The pathophysiological changes evaluated using this system initially designed for assessing the severity of a sepsis mouse model are broadly applicable to other mouse models of inflammatory disorders and measuring disease progression. Mice were initially monitored and scored prior to treatment and approximately 6h later. On the next day mice were again weighted and scored prior to treatment and about 6h post treatment. Then, scoring and body weigh were performed every morning until the end of the 10-day monitoring.

### Brain processing, IBA1 immunohistochemistry, and quantification

Brains were excised and passively perfused in 4% PFA solution at 4℃ for 72h and then transferred to 30% sucrose solution. Following embedding in OCT, sagittal brain sections (10 μm) were prepared on a Leica CM1850 cryostat. Following fixation, frozen sections were permeabilized with 0.1% Triton X-100 PBS solution and incubated in blocking solution consisting of 10% Normal donkey serum (Southern Biotech, 0030-01) and 0.1% Triton X-100 in 1X PBS for one hour at room temperature. Next, tissues were incubated with rabbit Anti-IBA1 (FUJIFILM Wako Pure Chemical Corporation, 019-19741) diluted in 0.1% Triton-100 and 10% NDS with 1X PBS overnight at 4℃. Next, sections were rinsed three times with 1x PBS-Triton X-100 (0.1M PBS and 0.1% Triton X-100) and incubated with donkey Anti-Rabbit Alexa Fluor 647 (ThermoFisher, A32795) diluted in 0.1% Triton-100 and 10% NDS with 1X PBS overnight at 4℃. Lastly, tissues were rinsed with 0.1% Triton X-100 in 1X PBS and counterstained with DAPI (Thermo Scientific, 62248). Slides were cover slipped with Fluoromount Mounting Medium (SouthernBiotech, 0100-01). Fluorescent images were captured with a Zeiss LSM 900 confocal microscope. Images were processed and analyzed using ZEISS ZEN blue software. Microglia density was analyzed by a wide count of IBA1+ cells across the hippocampal region. IBA1+ cells were selected for evident expression of DAPI+ nuclear staining with cytoplasmic staining of IBA1 (**Supplementary Figure 4**). Following the count, microglia cells were selected within the CA1 region of the hippocampus to assess the degree of ramification. Utilizing a previously established ramification index, individual IBA1+ cells were first marked for their cell body area, which encompasses the branching and extension of microglial processes (include citation number here). The perimeter of these microglial processes was then marked to assess the area covered by the total length of these processes. The ramification index was calculated by dividing the area by the perimeter, with a lower ratio of cell body area to the perimeter reflecting a more ramified microglia morphology.

### Statistics

Data are presented as means ± standard error of the mean. Statistical analyses of experimental data were conducted using GraphPad Prism 9.5.0 (GraphPad Software, Inc., La Jolla, CA). After evaluating the data for normality using a Shapiro-Wilk test, differences between groups were assessed using one-way ANOVA or the Kruskal Wallis test with appropriate post hoc test for multiple comparisons. Typically, data was pooled from experiments with multiple small cohorts of animals. Functional assessment curves and body weight were analyzed using repeated measures two-way ANOVA. All individual points refer to one animal. P values less than 0.05 were considered significant. All specific statistical tests are listed in the figure legends.

## Acknowledgements

This work was supported by the National Institutes of Health (NIH), National Institute of General Medical Sciences grants: RO1GM128008 and RO1GM121102 (to VAP), and R35GM118182 (to KJT).

## Data availability statement

The datasets generated during the current study are available from the corresponding author on reasonable request.

## Authors’ contributions

AF and VAP conceived the project. AF, VAP, AT, CNM, and EHC designed experiments. AF, SC, and SPP performed experiments. AF, SC, EHC, and VAP analyzed data. All authors discussed the results and commented on the manuscript. VAP wrote the first draft and AF, MB, EHC, SSC, FCC, AT, CNM, FCC, and KJT provided comments. All authors reviewed the manuscript and provided additional comments. All authors read and approved the final version of the manuscript.

---

**Supplementary Figure 1.**
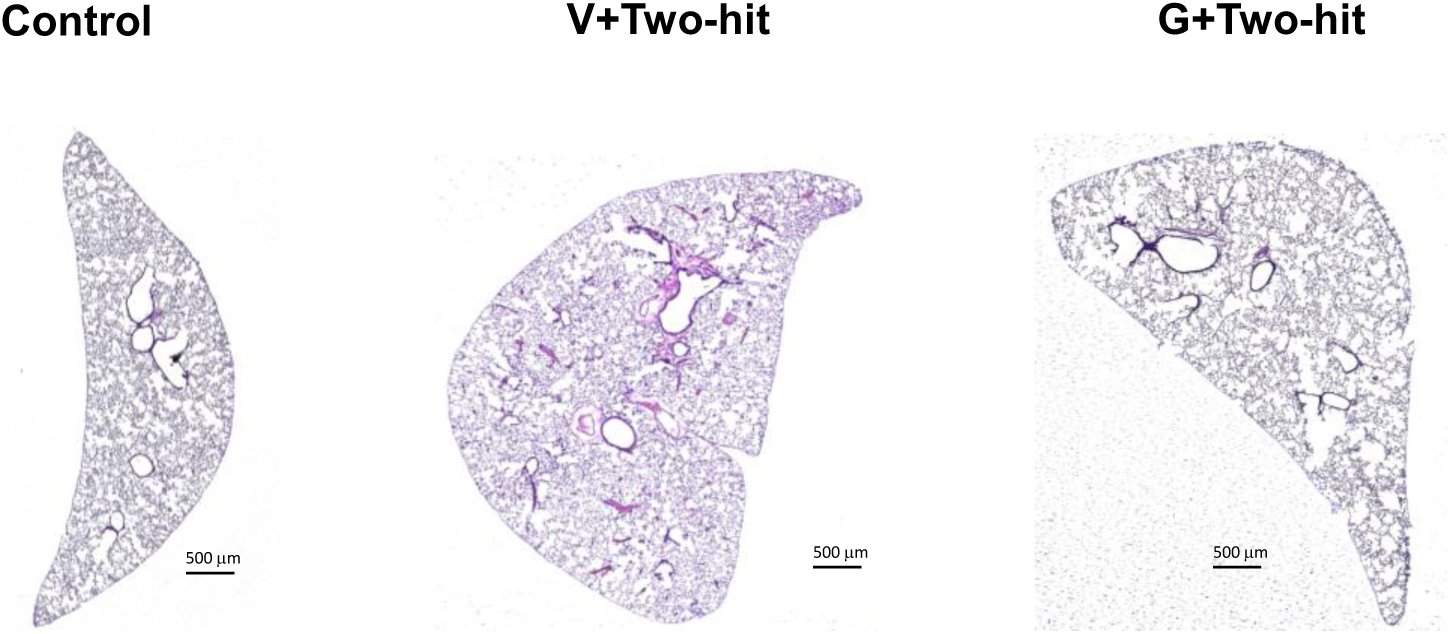
Whole lung images of control mice and mice subjected to ALI/ARDS and treated with vehicle or galantamine at 30h. Lung tissue was processed for H&E staining as described in Materials and Methods.

**Supplementary Figure 2.**
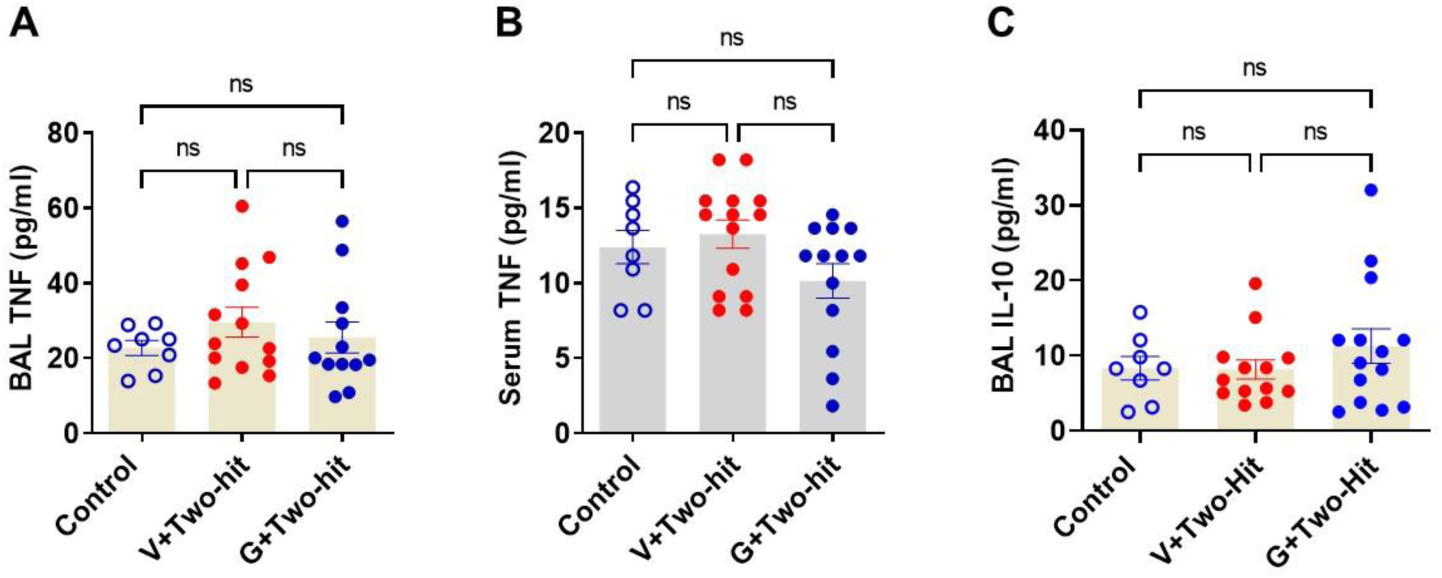
At 10 days BAL and serum TNF, and BAL IL-10 levels in ALI/ARDS mice treated with vehicle are not statistically different compared with control mice and ALI/ARDS mice treated with galantamine.

**Supplementary Figure 3.**
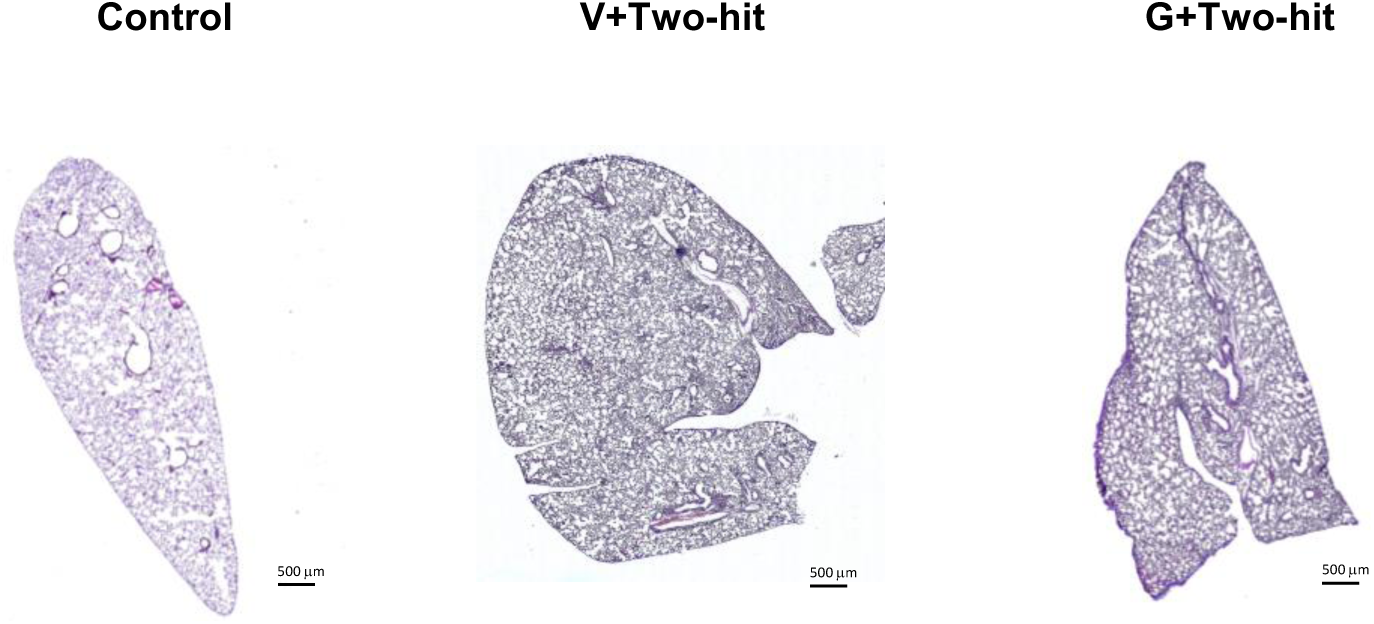
Whole lung images of control mice and mice subjected to ALI/ARDS and treated with vehicle or galantamine at 10 days. Lung tissue was processed for H&E staining as described in Materials and Methods.

**Supplementary Figure 4.**
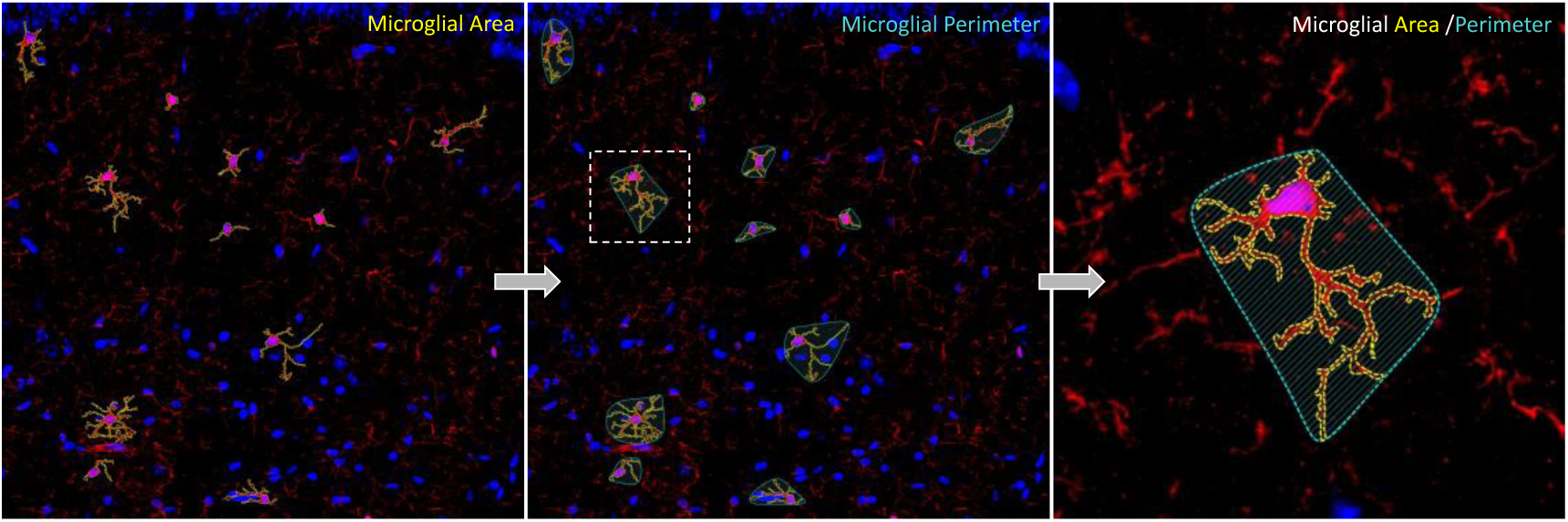
Detailed image of ramification index analysis taking place in the CA1 region of the hippocampus with IBA1+ shown in red and DAPI in blue. Yellow annotation marks the extent of microglial projections (area). Blue annotation marks total distance (perimeter) of projections that encompass single microglia.

